# Poly-ubiquitylated transmembrane proteins outcompete other cargo for limited space inside clathrin-coated vesicles

**DOI:** 10.1101/2024.12.17.628947

**Authors:** Hao-Yang Liu, Grant Ashby, Feng Yuan, Susovan Sarkar, Carl C. Hayden, Jon M. Huibregtse, Jeanne C. Stachowiak

## Abstract

Endocytic recycling of transmembrane proteins is essential to cell signaling, ligand uptake, protein traffic and degradation. The intracellular domains of many transmembrane proteins are ubiquitylated, which promotes their internalization by clathrin-mediated endocytosis. How might this enhanced internalization impact endocytic uptake of transmembrane proteins that lack ubiquitylation? Recent work demonstrates that diverse transmembrane proteins compete for space within highly crowded endocytic structures, suggesting that enhanced internalization of one group of transmembrane proteins may come at the expense of other groups. Here we show that preferential internalization of poly-ubiquitylated transmembrane proteins results in reduced endocytosis of mono-ubiquitylated and non-ubiquitylated proteins. Using a combination of live-cell imaging and ligand uptake assays, we confirmed that increased ubiquitylation correlates with increased internalization by clathrin-coated vesicles. Further, poly-ubiquitylated receptors significantly outcompeted their mono-ubiquitylated and non-ubiquitylated counterparts for localization to endocytic sites and uptake of extracellular ligands. These findings demonstrate the inherent interdependence of transmembrane protein recycling, suggesting that clathrin-coated vesicles act as selective filters, prioritizing highly ubiquitylated transmembrane proteins for uptake while leaving proteins with little or no ubiquitylation behind. Given that poly-ubiquitylation is thought to signal protein aging and damage, our findings suggest a mechanism for selective internalization of high priority cargo proteins, with simultaneously exclusion and protection of functional proteins that lack poly-ubiquitylation.

**Significance:** Ubiquitylation is essential for maintaining cellular homeostasis by regulating transmembrane protein trafficking, degradation, and signaling. Traditionally, poly-ubiquitylation has been viewed primarily as a signal for protein degradation, while mono-ubiquitylation is considered sufficient to trigger endocytosis. However, our work reveals a previously unrecognized role for poly-ubiquitylation in clathrin-mediated endocytosis, demonstrating that poly-ubiquitylated transmembrane proteins outcompete their non-ubiquitylated counterparts for incorporation into clathrin-coated vesicles, thereby establishing a competitive framework for endocytic cargo sorting. This mechanism reveals a selective sorting mechanism driven by the extent of ubiquitylation, which could regulate the removal of damaged proteins while protecting functional proteins at the plasma membrane.

## Introduction

Recycling of transmembrane proteins by clathrin-mediated endocytosis (CME) is crucial for maintaining homeostasis of the plasma membrane and sensitivity to the cellular environment through repeated cycles of ligand binding and signal transduction (1–4). During endocytic uptake, transmembrane proteins are first recognized as cargo by endocytic adaptor proteins, such as AP2, which subsequently recruit clathrin to form clathrin-coated vesicles that encapsulate the cargo (5–7). As the vesicles mature, additional transmembrane proteins and adaptor proteins are incorporated. Once the vesicle is fully formed, scission proteins such as dynamin facilitate the separation of the vesicle from the plasma membrane, allowing it to enter the cytoplasm for further processing (8).

Ubiquitylation of transmembrane proteins has long been known to increase their internalization by clathrin-coated vesicles (9–12). This post-translational modification, which involves the attachment of ubiquitin to lysine residues within the intracellular domains of transmembrane proteins, enhances their affinity for key components of clathrin-coated vesicles (10, 11). Specifically, Eps15, a protein that arrives early in the assembly of clathrin-coated vesicles, recruits ubiquitylated cargo using its tandem ubiquitin-interacting motifs (UIMs) (13–15). These interactions stabilize the initiation of clathrin-coated vesicles, resulting in an increase in the fraction of endocytic assemblies that successfully internalize cargo (16). Similarly, Epsin, a key component of mature clathrin-coated vesicles, contains UIMs which enable it to function as an adaptor for ubiquitylated cargo proteins (17–21). Interestingly, the UIMs of Epsin1 exhibit a selectively binding affinity for poly-ubiquitin chains, while showing minimal interaction with mono-ubiquitin. These observations suggest that endocytic sites may preferentially recognize poly-ubiquitin chains as sorting signals (22, 23).

However, previous work suggests that both mono-ubiquitylation and poly-ubiquitylation of transmembrane proteins can promote their uptake by clathrin-mediated endocytosis. Some studies in yeast cells indicate that mono-ubiquitylation efficiently directs surface proteins to cortical actin patches, the sites of clathrin-mediated uptake in *S. cerevisiae,* while poly-ubiquitin chains signal proteasomal degradation (24–26). However, the Fur4 transmembrane protein, which is predominately poly-ubiquitylated, is also internalized by clathrin-mediated endocytosis in yeast cells, suggesting a lack of clarity in the role of mono versus poly-ubiquitin during endocytosis in yeast (27, 28). Similarly, in mammalian cells, the role of ubiquitylation during internalization of the epidermal growth factor receptor (EGFR) remains unclear. Early studies suggested that EGFR was exclusively mono-ubiquitylated. In contrast, more recent findings reveal that within five minutes of activation, EGFR is predominantly poly-ubiquitylated, creating uncertainty about which form best promotes endocytosis (29, 30).

Importantly, all of these studies examine endocytosis of individual transmembrane proteins in isolation from the other proteins at the plasma membrane surface. However, recent work from our lab and others suggest that endocytic structures are highly crowded, such that transmembrane proteins must compete with one another for limited space inside endocytic carriers (31–35). In particular, bulky glycosylated transmembrane proteins, which take up more space inside of endocytic structures, and proteins with lower affinity for the endocytic machinery were outcompeted by their smaller, higher affinity counterparts during endocytic internalization (31, 36). In this competitive landscape, how might the enhanced uptake of ubiquitylated transmembrane proteins impact internalization of proteins with little or no ubiquitylation?

To examine this question, we performed a series of experiments using genetically engineered transmembrane proteins with varying degrees of stable ubiquitylation. We examined the localization of these model transmembrane proteins to endocytic sites and measured their ability to internalize ligands. Our findings demonstrate that poly-ubiquitylated proteins not only localize more robustly to endocytic structures but also exhibit higher internalization efficiency compared to their mono-ubiquitylated and non-ubiquitylated counterparts. This observation suggests that poly-ubiquitylated transmembrane proteins may be prioritized during internalization. To investigate this possibility, we developed assays in which transmembrane proteins with varying degrees of stable ubiquitylation competed for localization to endocytic sites and for ligand uptake. Our results demonstrate that poly-ubiquitylated transmembrane proteins consistently outcompeted their mono-ubiquitylated and non-ubiquitylated counterparts, indicating a selective advantage during endocytic uptake. Together, our findings suggest that clathrin-mediated endocytosis is highly sensitive to the extent of ubiquitylation of transmembrane proteins, resulting in selective uptake of poly-ubiquitylated cargo. Given that damaged protein and protein that have completed their function in cell signaling are frequent targets of poly-ubiquitylation (29, 37–40), these findings could help to explain how endocytic structures select key populations of transmembrane proteins for rapid removal from the cell surface, leaving functional transmembrane proteins behind.

## Results

### Poly-ubiquitylation increases endocytic uptake of model transmembrane proteins relative to mono-ubiquitylation

To evaluate the role of ubiquitylation in the partitioning of transmembrane proteins to endocytic sites, we designed chimeric membrane proteins with a defined number of enzymatically inert ubiquitin moieties (0, 1, or 4) fused to the intracellular and transmembrane domains of the transferrin receptor (TfR), followed by an extracellular red fluorescent protein (RFP) domain for visualization (Fig. A-C). TfR was selected as the model protein due to its well-characterized, strong constitutive internalization via clathrin-mediated endocytosis (CME) (41–44). These model transmembrane proteins will henceforth be referred to as 0-Ub, 1-Ub, and 4-Ub. The native intracellular domain of TfR contains a YTRF motif, which mediates constitutive internalization at endocytic sites by interacting with the μ2 subunit of the clathrin adaptor protein AP2 (45–47). During internalization, TfR may undergo mono-ubiquitylation mediated by MARCH8, whose N-terminal RING-CH domain specifically recognizes the transmembrane region of TfR and catalyzes ubiquitylation of its cytoplasmic domain (48). To prevent endogenous ubiquitylation by cellular ligases, we mutated all four lysine residues in the TfR intracellular domain and all seven lysines within the ubiquitin moieties to arginine. Additionally, to maintain the stability of attached ubiquitin moieties, the C-terminal GG residues of all ubiquitin moieties were mutated to AA to prevent deubiquitylation. Further, the chimeric construct lacked the extracellular domain that normally mediates TfR dimerization, resulting in a monomeric form and eliminating potential dimerization-related influences on the role of ubiquitylation.

Each of the model transmembrane proteins was separately expressed in SUM159 human breast cancer-derived epithelial cells. We chose these cells because of their thin lamellipodia, which are ideal for visualization of endocytic events. These cells were gene-edited to incorporate a HaloTag on both alleles of the σ2 subunit of AP2, the major adaptor protein of the clathrin pathway (49, 50). Notably, AP2, which is thought to be in approximately 1:1 stoichiometric ratio with clathrin, is preferred over clathrin for tracking assembly of endocytic vesicles, because AP2 localizes exclusively to the plasma membrane, while clathrin is also found at endosomes and other organelles (51). For visualization of AP2 during fluorescence imaging, we used the far-red HaloTag ligand, JF_646_ (52). We imaged the plasma membrane of adherent cells using spinning disk confocal microscopy (*SI Appendix*, Fig. S1). First, we examined cells expressing the 0-Ub model protein, which exhibited clear colocalization of the transmembrane fusion protein (RFP) with endocytic sites (AP2), suggesting robust internalization of the model protein as expected for the transferrin intracellular domain (Fig. 1D). Next, we examined confocal images of cells expressing the 1-Ub and 4-Ub model proteins. As the number of ubiquitins increased, the intensity of the model proteins (RFP) that colocalized with puncta in the AP2 channel (JF_646_) appeared to increase (Fig. 1E, F).

**Figure 1.**
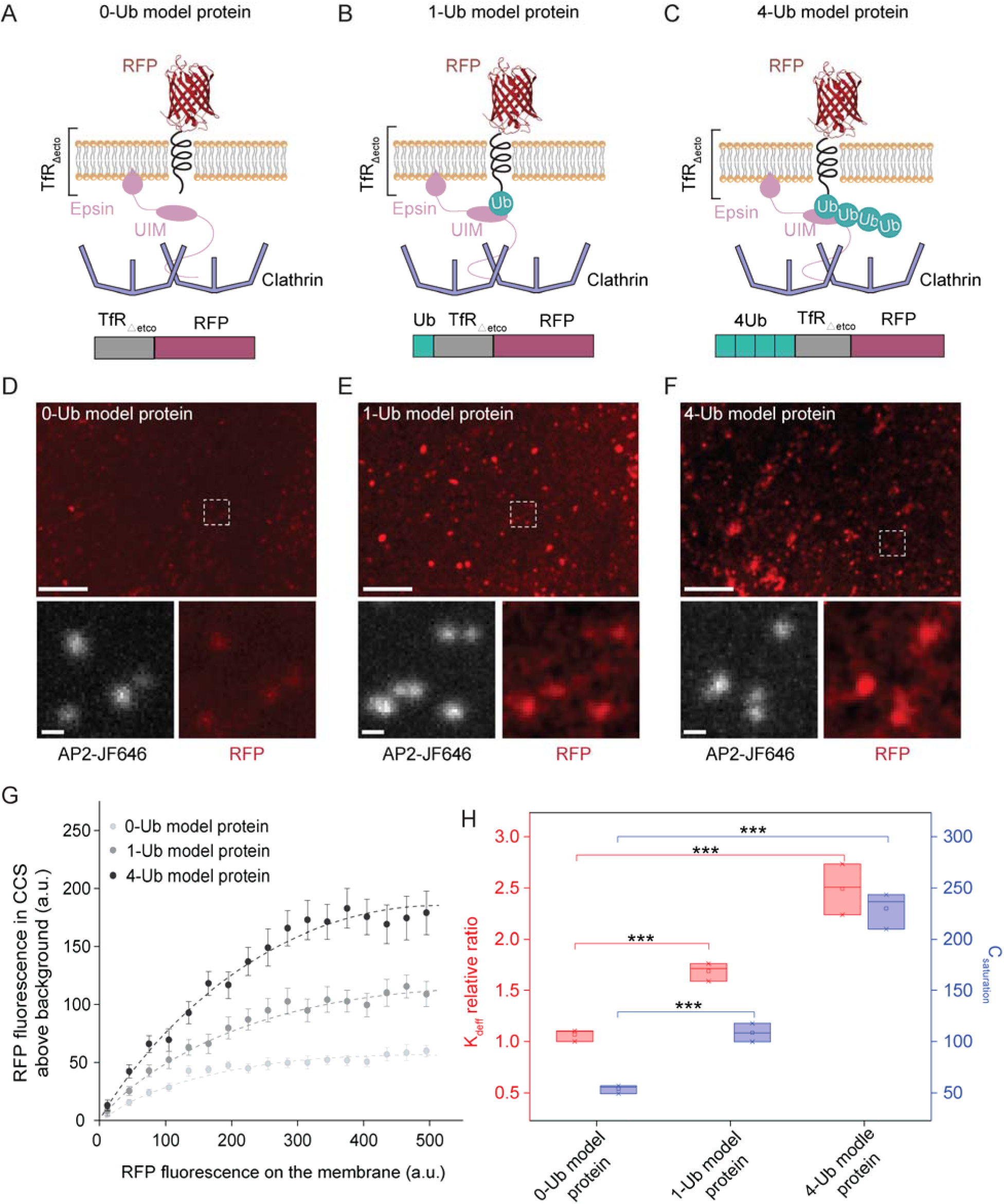
Poly-ubiquitylation increases endocytic uptake of model transmembrane proteins relative to mono-ubiquitylation. (A-C) Schematic representations of transmembrane fusion proteins with defined ubiquitin modifications: (A) the 0-Ub model protein with no attached ubiquitin, (B) the 1-Ub model protein with a single attached ubiquitin, and (C) the 4-Ub model protein with four attached ubiquitins. All lysine residues in the cytoplasmic domain of TfR were mutated to arginines to prevent ubiquitylation. To further stabilize the attached ubiquitin moieties all lysines within ubiquitin were mutated to arginines, and the C-terminal GG residues were mutated to AA to prevent intracellular deubiquitylation. Epsin interacts with ubiquitylated cargo through its ubiquitin-interacting motif (UIM) and facilitates clathrin-mediated endocytosis by binding to clathrin. (D-F) Spinning disk confocal images of the plasma membrane of SUM 159 cells transiently expressing the 0-Ub model protein (D), 1-Ub model protein (E) and 4-Ub model protein (F). The white boxes in the upper panels indicate regions magnified in the lower insets. Red fluorescence (RFP) highlights the model proteins, while white fluorescence (AP2-JF646) marks endocytic sites. The scale bars are 5 µm (main image) and 500 nm (insets). (G). The relative number of transmembrane fusion proteins localized in clathrin-coated structures is shown against the relative concentration of fusion proteins on the plasma membrane around each structure. Data were collected from 11196 endocytic sites in 80 cells expressing 0-Ub model protein, 12361 endocytic sites in 85 cells expressing 1-Ub model protein, and 8732 endocytic sites in 68 cells expressing 4-Ub model protein. Each point reflects the average value from clathrin-coated structures binned by the local membrane concentration of the proteins. Error bars denote mean ± s.e., while dashed lines represent model fits based on the best-fit values of Kdeff and Nmax. (H). Box plot showing the relative Kdeff ratio values (red, left Y-axis) and Csaturation values (blue, right Y-axis) for the 0-Ub, 1-Ub, and 4-Ub model proteins. Kdeff represents the effective binding constant, normalized to the 0-Ub model protein, while Csaturation represents the maximum local concentration of fusion proteins in clathrin-coated structures. For each box, n=3 biologically independent experiments with at least 40 cells for each condition. A two-sample *t* test was conducted for pairwise comparisons. P-values were <0.05 between each pair of fusion proteins suggesting a statistically significant difference. **P < 0.01, ***P < 0.001.

We quantified the partitioning of each model transmembrane protein to endocytic sites using a publicly available algorithm, CMEAnalysis (53). This algorithm applies a 2D Gaussian fit to puncta in the AP2 channel, which was designated as the “master channel” to identify endocytic structures, and subsequently to the corresponding locations in the transmembrane fusion protein channel, which served as the “subordinate channel” for estimating the intensity of each structure relative to the local plasma membrane background signal. The amplitudes of these Gaussian fits report the relative number of model proteins per endocytic structure (54). The algorithm also records the average fluorescence intensity of model proteins on the plasma membrane in the immediate vicinity of each punctate structure. This intensity represents the relative concentration of model proteins on the surrounding plasma membrane (54). Using these data, we plotted the relative number of model proteins per endocytic structure as a function of the relative concentration of model proteins in the surrounding plasma membrane. The resulting plots showed that the relative number of model proteins within each clathrin-coated structure initially increased with increasing concentration of model proteins in the surrounding plasma membrane. However, as the concentration on the plasma membrane further increased, the relative number of model proteins in endocytic sites began to plateau toward a maximum value. This maximum reflects the relative number of transmembrane model proteins necessary to fully saturate a clathrin-coated structure, as previously described (54), Fig. 1G.

These data suggest that the saturated capacity of clathrin-coated structures for model transmembrane proteins significantly increased as the number of ubiquitins increased, from zero to four. To estimate the saturation value, we used a simple multivalent binding model that we have described previously (Equation 1) (54). Here the average number of model proteins per structure, denoted as ⟨*n*⟩, is determined by the maximum capacity per endocytic structure (N_max_), the relative concentration of fusion proteins on the plasma membrane (C_mem_), and the dissociation constant (K_deff_) representing the affinity between the fusion protein and the endocytic structure.

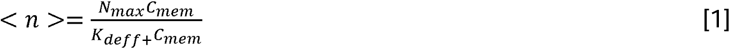

This equation assumes that the number of model proteins per endocytic structure is governed by a dynamic balance between model proteins inside and outside the structure. We applied this equation to the data in Fig. 1G using N_max_ and K_deff_ as free parameters. Fitting this model to the data suggests the relative affinity (K_deff_) of the model proteins for endocytic sites increased with the number of ubiquitins conjugated to it. Specifically, K_deff_ increased by 2.3-fold for 4-Ub compared to 0-Ub, and by 1.6-fold for 1-Ub compared to 0-Ub (Fig. 1H). This result is expected, as multivalent interactions between endocytic proteins and ubiquitin likely increase the overall affinity between the model protein and endocytic structures. What was surprising was that the saturated capacity of the endocytic structures, N_max_, also increased, 3.3-fold for 4-Ub compared to 0-Ub and 2.1-fold for 1-Ub compared to 0-Ub. (Fig. 1H). This finding suggests that endocytic sites are able to accommodate a substantially larger number of poly-ubiquitylated model proteins in comparison to mono-ubiquitylated or non-ubiquitylated proteins. If so, then a similar trend should exist for endocytic internalization of the model proteins. Therefore, we next sought to evaluate internalization of model proteins using flow cytometry-based assays.

### Poly-ubiquitin chain length controls ligand uptake by model receptors

We next evaluated to what extent the impact of ubiquitin on localization to endocytic sites resulted in an increase in ligand/receptor uptake by SUM159 cells. To this end, we designed a new model receptor-ligand system, as shown in Fig. 2 A-C. Similar to the model transmembrane proteins in Fig. 1, the N-terminus of the model receptors begins with a fixed number of stable ubiquitins (0, 1, or 4), followed by the TfR intracellular and transmembrane domains. The extracellular domain consists of an RFP domain for visualization followed by a C-terminal domain consisting of a single-domain camelid antibody (i.e., nanobody) with nanomolar affinity for GFP (55). As before, the lysines in the intracellular domain of TfR and the linked ubiquitins were mutated to arginines. We transiently expressed each model receptor in SUM159 cells, which were subsequently incubated with a solution of 500 nM GFP for one hour. Confocal images of these cells revealed clear colocalization between the model receptor (RFP), its ligand (GFP), and endocytic sites (AP2-JF_646_). The intensity of the receptor and ligand at endocytic sites appeared to increase with increased length of the ubiquitin chain (0, 1, 4), as expected (Fig. 2 A-C).

**Figure. 2.**
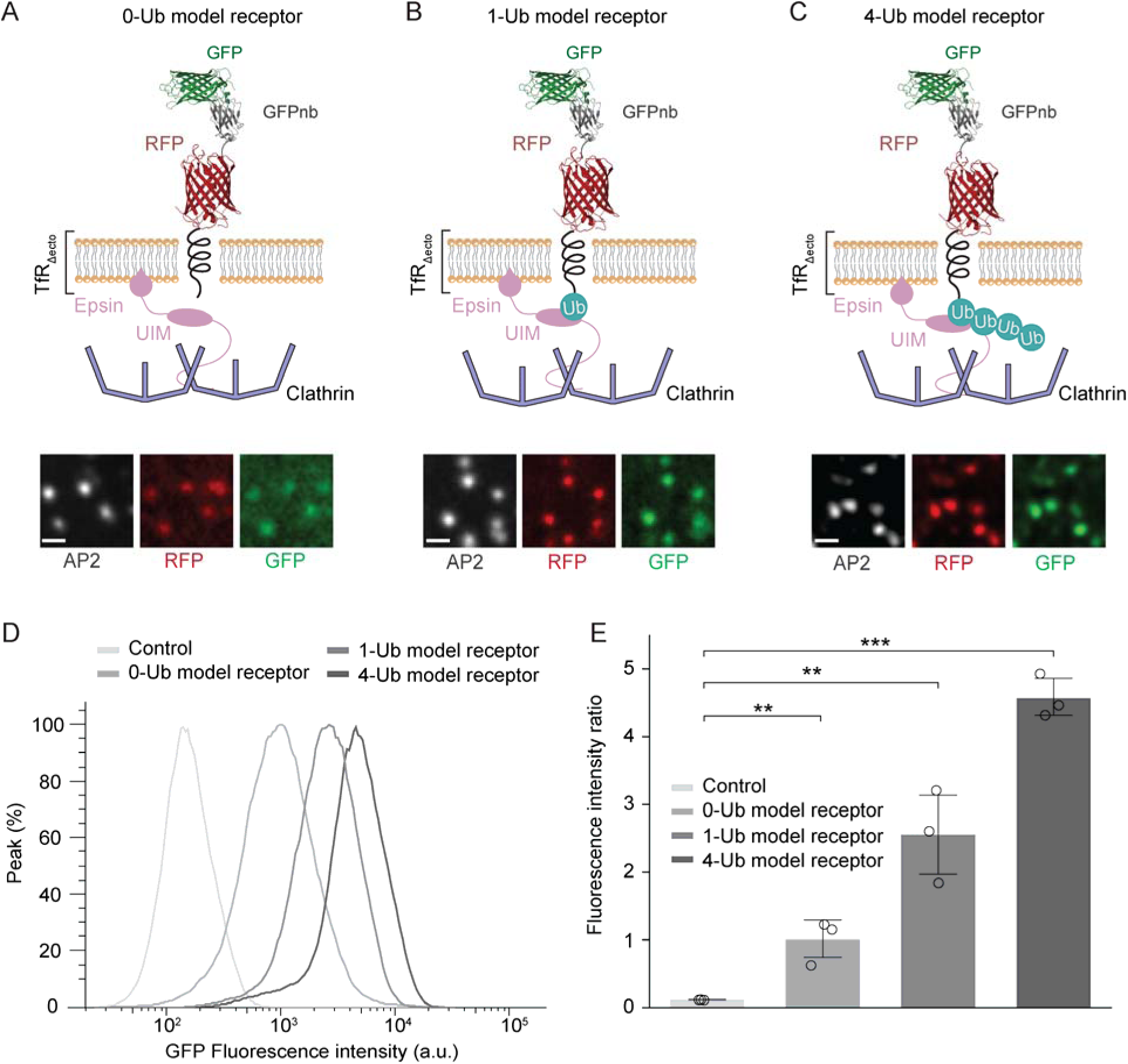
Poly-ubiquitin chain length controls ligand uptake by model receptors. (A-C) Schematic illustrations of model receptors with defined number of ubiquitins bound to GFP: (A) 0-Ub model receptor, (B) 1-Ub model receptor and (C) 4-Ub model receptor. All lysine residues in the cytoplasmic domain of TfR were substituted with arginines to inhibit ubiquitylation. Similarly, lysine residues within ubiquitin were replaced with arginines, and the C-terminal GG residues were mutated to AA to prevent intracellular deubiquitylation. Fluorescence images in the lower panel display the plasma membrane of SUM159 cells expressing each model receptor, showing the colocalization of AP2 (white), RFP (red), and GFP (green). Scale bar: 1 μm. (D) Representative histograms from flow cytometry demonstrating GFP uptake in SUM159 cells expressing the model receptors, with increased uptake corresponding to higher levels of ubiquitylation. (E) Quantification of GFP uptake in SUM159 cells expressing 0-Ub, 1-Ub, or 4-Ub model receptors, as well as a control, indicating a positive correlation between poly-ubiquitin chain length and receptor internalization. n=3 biologically independent samples for each group. Data are mean ± s.e.. Two-sample t-tests showed statistically significant differences between groups (**p < 0.01, ***p < 0.001).

After incubation with GFP, the cells were trypsinized and thoroughly washed to ensure removal of unbound GFP from the cell surface and the culture medium (56). The total GFP uptake for each cell population was then quantified using flow cytometry (*SI Appendix*, Fig. S2 and S3). As shown in Fig. 2D, the intensity distribution of GFP fluorescence shifted toward higher values for cells expressing the 0-Ub model receptor, compared to the untransfected control group, indicating uptake of GFP. Comparing the distributions for the three model receptors (Fig. 2E), cells expressing the model receptor with 1-Ub showed a 2.6-fold increase in GFP uptake compared to cells expressing the 0-Ub model receptor, while cells expressing the 4-Ub receptor showed a 4.8-fold increase compared to the 0-Ub group. These results confirm that the increase in localization to endocytic sites observed for ubiquitylated receptors in Fig. 1 correlates with an increase of similar magnitude in their internalization by cells.

### Poly-ubiquitylation of model receptors increases stability of endocytic structures relative to mono-ubiquitylation

The increase in the number of ubiquitylated model proteins required to saturate endocytic structures (Fig. 1G, H), which correlates with an increase in ligand uptake by ubiquitylated model receptors (Fig. 2D, E), suggests that clathrin-coated vesicles have a larger capacity for ubiquitylated transmembrane proteins compared to their non-ubiquitylated counterparts. How is this increase in capacity achieved? We considered three nonexclusive explanations (Fig. 3A). First, given that ubiquitin plays an important role in stabilizing endocytic sites (16, 57), the presence of a significant number of ubiquitylated transmembrane proteins at the plasma membrane could cause endocytic sites to grow physically larger such that they can accommodate a larger amount of transmembrane protein cargo. Second, the increased stability of endocytic sites in the presence of ubiquitylated transmembrane proteins could result in a greater fraction of endocytic sites successfully maturing into productive endocytic vesicles that internalize cargo. Third, the increased affinity of ubiquitylated transmembrane proteins for endocytic sites could enable these proteins to effectively outcompete other transmembrane proteins for limited space within endocytic structures with a fixed capacity.

**Figure 3.**
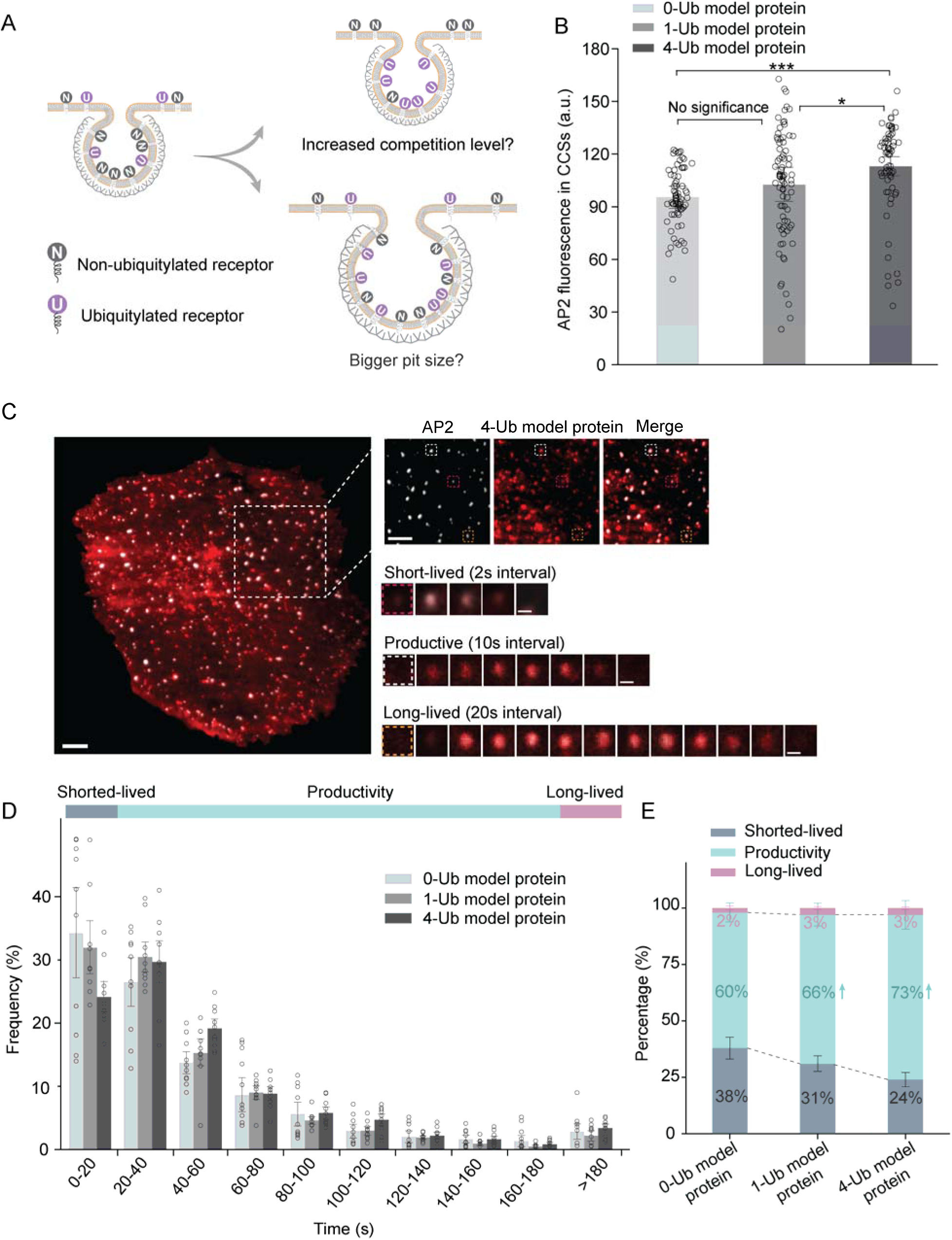
Poly-ubiquitylation of model receptors increases the stability of endocytic structures relative to mono-ubiquitylation. (A) Schematic illustration of two possible mechanisms by which receptor ubiquitylation impacts clathrin-coated vesicle formation: (i) poly-ubiquitylation increases the size of clathrin-coated pits, leading to larger vesicles, or (ii) poly-ubiquitylation enhances the competition among ubiquitylated receptors and other cargos for limited space within the vesicles. (B) Quantification of AP2 fluorescence intensity in CCSs from SUM159 cells expressing 0-Ub, 1-Ub, or 4-Ub model proteins. As AP2 and clathrin exhibit a nearly 1:1 stoichiometric ratio, the relative abundance of AP2 molecules per structure can be used to estimate the size of endocytic structures. AP2 fluorescence increases with the increased ubiquitylation levels of the receptors, indicating a bigger size of CCS. n=25 biologically independent cell samples for each group. A two-sample *t* test was conducted for pairwise comparisons. P-values were <0.05 between each group suggesting a statistically significant difference, ***P < 0.001. (C) Representative TIRF image of a SUM159 cell expressing 4-Ub model protein: AP2 (white) and 4-Ub model protein (red). Large inset highlights three representative clathrin-coated structures shown in smaller insets: short-lived (red), productive (white), and long-lived (orange) structures lasting 8 s, 70 s, and 240 s, respectively. Scale bars: 5 µm (main image and magnified panels on the right), 500 nm (time series images). (D) Histograms of lifetime distributions of clathrin-coated structures for each group. Data were collected from n=10 biologically independent cell samples. Structures with lifetimes shorter than 20 s are categorized as short-lived, those between 20 and 180 s are labeled as productive, and structures lasting longer than 180 s are classified as long-lived. 0-Ub model protein n=15 biologically independent cell samples, 15120 pits; 1-Ub model protein n=12 biologically independent cell samples, 13790 pits; 0-Ub model protein n=11 biologically independent cell samples, 7350 pits; (E) Stacked column plot shows the proportion of clathrin-coated structures categorized as short-lived (dark blue), productive (light blue), and long-lived (pink) for 0-Ub, 1-Ub and 4-Ub model proteins, respectively. Data are mean ± s.e.. An upward arrow (↑) denotes a significant increase at the 95 % confidence level compared to the 0-Ub model protein group.

First, we assessed the impact of transmembrane protein ubiquitylation on the size of clathrin-coated vesicles. Using CMEAnalysis, we quantified the intensity of endocytic structures in the AP2 channel (JF_646_) for the model transmembrane proteins containing varying numbers of ubiquitins. Given the near 1:1 stoichiometric ratio between AP2 and clathrin, the relative number of AP2 proteins per structure can serve as a rough proxy for the size of endocytic structures (51). Fig. 3B shows the average AP2 fluorescence intensity for endocytic structures in cells expressing each of the model transmembrane proteins (0-Ub, 1-Ub, 4-Ub). These data suggest modest increases in the size of endocytic structures for cells expressing the 1-Ub (7% brighter) and 4-Ub (18% brighter) model transmembrane proteins, compared to cells expressing the 0-Ub model transmembrane protein.

Next, we examined the impact of the 0-Ub, 1-Ub and 4-Ub model transmembrane proteins on the stability of growing endocytic structures. For this purpose, total internal reflection fluorescence (TIRF) microscopy was used to image the recruitment of model proteins into growing endocytic structures in real time (*SI Appendix*, Fig. S4). TIRF is ideal for tracking the dynamics of endocytosis due to the shallow penetration depth (∼100 nm) of the evanescent field, which selectively illuminates the plasma membrane while minimizing fluorescence from the cytoplasm and organelles. We captured images every 2s, collecting 250 frames in total across the same two fluorescent channels used above: (i) AP2 (JF_646_) and (ii) transmembrane model proteins (RFP).

The time series were analyzed to track individual endocytic events from initiation to departure from the TIRF field. The lifetime of an endocytic structure on the membrane, measured as the interval between the arrival and departure of a labeled marker, can range from seconds to minutes (58). Structures lasting less than 20 seconds are typically classified as ‘short-lived’ and are likely abortive, based on prior studies showing their inability to generate productive vesicles. In contrast, ‘productive’ structures, which successfully form vesicles, typically remain on the membrane for 20 seconds to several minutes. Structures that persist for over 180 seconds are considered ‘long-lived’ and are thought to have stalled, potentially undergoing delayed internalization or failing to form vesicles, Fig. 3C (59). Each of these populations was observed in our experiments, and the distribution of the lifetimes of endocytic structures for cells expressing each of the model transmembrane proteins is shown in Fig. 3D. The data indicate that the fraction of “short-lived” endocytic structures progressively decreased upon expression of the 1-Ub and 4-Ub model transmembrane proteins, which corresponded to an increase in the number of “productive” structures (10%, 22% for 1-Ub and 4-Ub, in comparison to 0-Ub, respectively), while the fraction of “long-lived” structures remained low and approximately constant, Fig. 3E. Taken together, data on the size and stability of endocytic structures indicate a significant increase upon expression of ubiquitylated transmembrane proteins.

### Poly-ubiquitylated model transmembrane proteins outcompete mono-ubiquitylated proteins for limited space within endocytic structures

We next evaluated the impact of ubiquitylation on competition between transmembrane proteins for limited space within clathrin-coated vesicles. Specifically, we co-expressed two model proteins simultaneously in SUM159 cells: one labeled with RFP and the other with GFP. The RFP-labeled model protein (TfR-RFP), which was not ubiquitylated, served as a “bystander”, while the GFP-labeled model protein served as a “competitor” and was conjugated with either 0, 1, or 4 ubiquitins, using the same domain structure as the model proteins in Fig. 1 (Fig. 4A-D). Confocal microscopy showed that both RFP and GFP-labeled model proteins localized to endocytic sites (AP2 puncta in JF_646_ channel). However, as the number of ubiquitins on the competitor increased, the partitioning of the RFP-labeled bystander into these sites decreased. Specifically, the contrast in the RFP channel between the endocytic sites and the surrounding plasma membrane weakened, while the GFP signal contrast progressively increased. These shifts suggest that the presence of a ubiquitylated competitor reduced the localization of the non-ubiquitylated bystander at endocytic sites. These trends are quantified in Fig. 4E, which indicates that the relative number of bystanders (RFP) per endocytic structure initially increased with the relative concentration of bystanders on the surrounding plasma membrane, before approaching a plateau corresponding to the saturated capacity of the endocytic structures for the bystander protein. Notably, as the GFP-tagged competitors are in close proximity to the RFP-tagged receptor on the plasma membrane, there is potential for fluorescence resonance energy transfer (FRET), which could reduce the intensity in the GFP channel. To avoid this potential artifact, we only analyzed the RFP channel images. Additionally, cells were selected based on their expression level of the GFP-tagged competitor, ensuring that variations in expression level were not misinterpreted as competition between the receptors.

**Figure 4.**
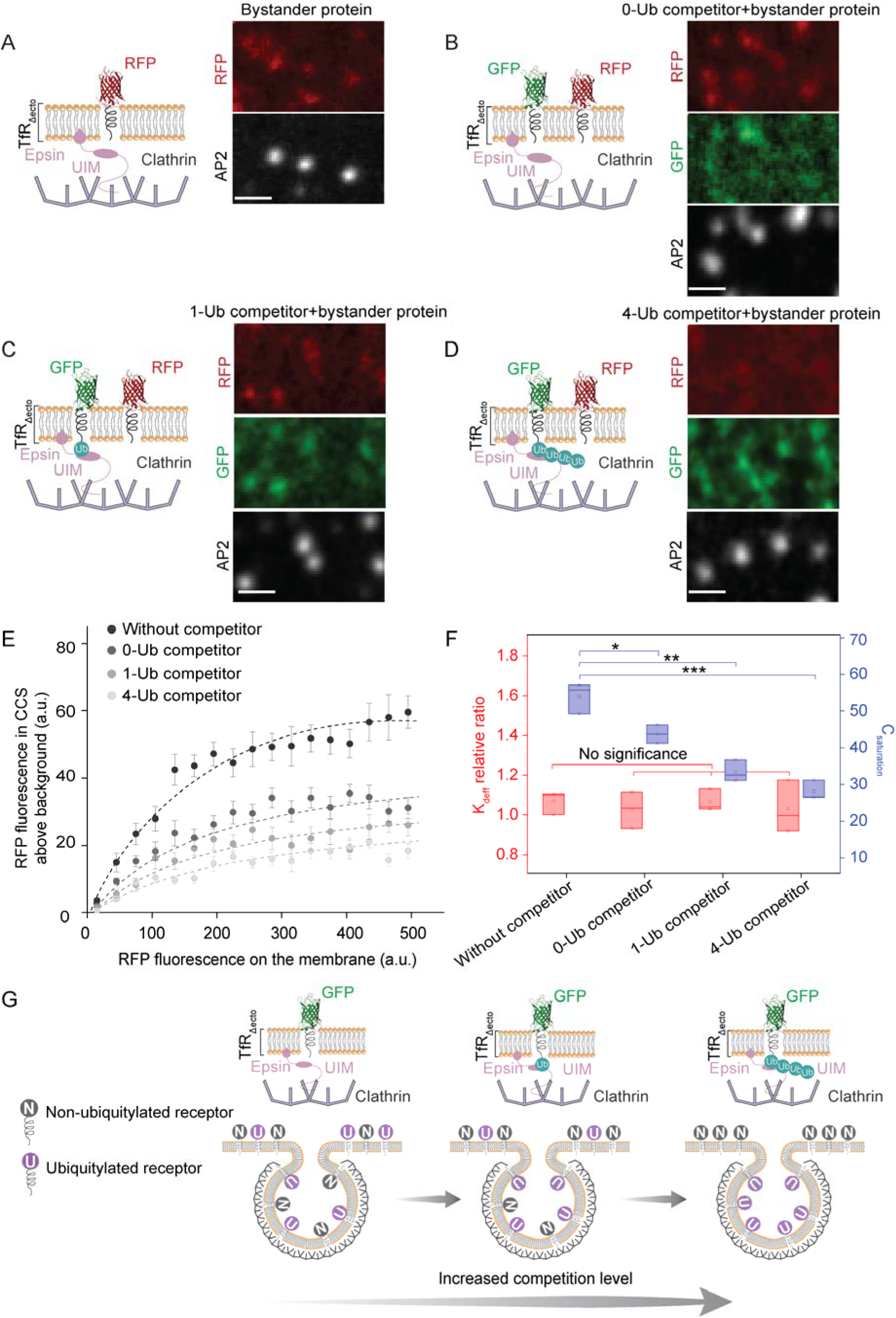
Poly-ubiquitylated model receptors outcompete mono-ubiquitylated model receptors for limited space within endocytic structures. (A–D) Schematic illustrations and fluorescence images depicting the incorporation of non-ubiquitylated bystander protein (RFP-labeled) into endocytic sites when competing with GFP-labeled competitors bearing different ubiquitin modifications: (A) without competitor, (B) with 0-Ub competitor, (C) with 1-Ub competitor, and (D) with 4-Ub competitor. All lysine residues in the cytoplasmic domain of TfR were mutated to arginines to prevent ubiquitylation. To stabilize the attached ubiquitin moieties, lysines within ubiquitin were mutated to arginines, and the C-terminal GG residues were altered to prevent intracellular deubiquitylation. Epsin binds to ubiquitylated cargo via its ubiquitin-interacting motif and promotes clathrin-mediated endocytosis by interacting with clathrin. Fluorescence images show red fluorescence (RFP) marking bystander proteins, green fluorescence (GFP) marking competitors, and white fluorescence (AP2-JF646) indicating endocytic sites. Dashed circles highlight areas of signal colocalization between RFP, GFP, and AP2, indicating the recruitment of bystander proteins and competitors to the same endocytic sites. Scale bar: 1 µm. (E) The relative number of bystander protein (RFP channel) localized in clathrin-coated structures is shown against the relative concentration of fusion proteins on the plasma membrane around each structure. Data were collected from 11196 endocytic sites in 80 cells expressing without competitor, 8038 endocytic sites in 60 cells expressing 0-Ub competitor, 8493 endocytic sites in 63 cells expressing 1-Ub competitor, and 11446 endocytic sites in 75 cells expressing 4-Ub competitor. Each point reflects the average value from clathrin-coated structures binned by the local membrane concentration of the proteins. GFP-labeled competing receptor expression is constrained to 200-300 GFP per μm^2^. Error bars denote mean ± s.e., while dashed lines represent model predictions based on the best-fit values of Kdeff and Nmax. (F) Box plot showing the relative Kdeff values (red, left Y-axis) and Csaturation values (blue, right Y-axis) for the control (without competitor), 0-Ub, 1-Ub, and 4-Ub competitors. Kdeff represents the effective binding constant, normalized to the 0-Ub model protein, while Csaturation represents the maximum local concentration of fusion proteins in clathrin-coated structures. Each box displays the median, interquartile range, and individual data points. A two-sample *t* test was conducted for pairwise comparisons. P-values were <0.05 between each group suggesting a statistically significant difference. **P < 0.01, ***P < 0.001. (G) Cartoon schematic illustrating the increased competition for endocytic sites between ubiquitylated receptors and other cargoes as the ubiquitylation level of competing receptors increases. The diagram shows how higher ubiquitylation levels enhance receptor recruitment to clathrin-coated vesicles, potentially leading to displacement of lower or normal (non-ubiquitylated) receptors.

The data clearly show that the saturated capacity of endocytic structures for the bystander protein (RFP labeled) is highest when the bystander is expressed alone, and progressively decreases when it is co-expressed alongside the 0-Ub or 4-Ub competitors (GFP labeled). Fitting these data with the multivalent binding model used above (Equation 1) provides measure of the impact of the competitor on the effective affinity of the bystander for endocytic sites (K_deff_), and the effective capacity of endocytic sites for the bystander (N_max_) (Fig. 4F). These trends indicate that the competitor had little impact on the affinity of the bystander for endocytic sites, as would be expected, given that the competitor is unlikely to alter the biochemistry of binding between the bystander and endocytic adaptor proteins such as AP2. In contrast, the presence of the competitor had a substantial influence on the effective capacity of endocytic sites for the bystander, lowering its saturation value by 1.2-fold for the 1-Ub competitor and 1.6-fold for the 4-Ub competitor, in comparison to the 0-Ub competitor. These results suggest that internalization of a transmembrane protein decreases substantially in the presence of more highly ubiquitylated competitors (Fig. 4G). Up to this point, our data suggest that the increased capacity of endocytic sites for ubiquitylated cargo can be explained by increases in both the stability of endocytic sites (Fig. 3D, E) and favorable competition with non-ubiquitylated cargo (Fig. 4), with a more minor contribution from an increase in the size of endocytic structures (Fig. 3B).

### Poly-ubiquitylated model receptors outcompete mono-ubiquitylated model receptors for ligand uptake

Next, we sought to further evaluate the competitive advantage of ubiquitylated transmembrane proteins by measuring their impact on ligand uptake during flow cytometry experiments. Specifically, we asked what impact a ubiquitylated “competitor” has on the ability of a non-ubiquitylated “bystander receptor” to internalize its ligand via endocytosis. For this purpose, we employed a non-ubiquitylated model receptor for GFP, identical to the one used in Fig. 2A, except that the RFP domain was replaced by BFP, leaving the RFP channel available for RFP-labeled competitors. Using flow cytometry, we measured the ability of the resulting “bystander receptor” to internalize GFP when expressed either alone, or in combination with RFP-tagged competitors having 0, 1, or 4 ubiquitins fused to their intracellular domains. As shown in Fig. 5A-D, these competitors did not bind to GFP such that the internalized GFP signal resulted entirely from internalization of the bystander receptors.

**Figure 5.**
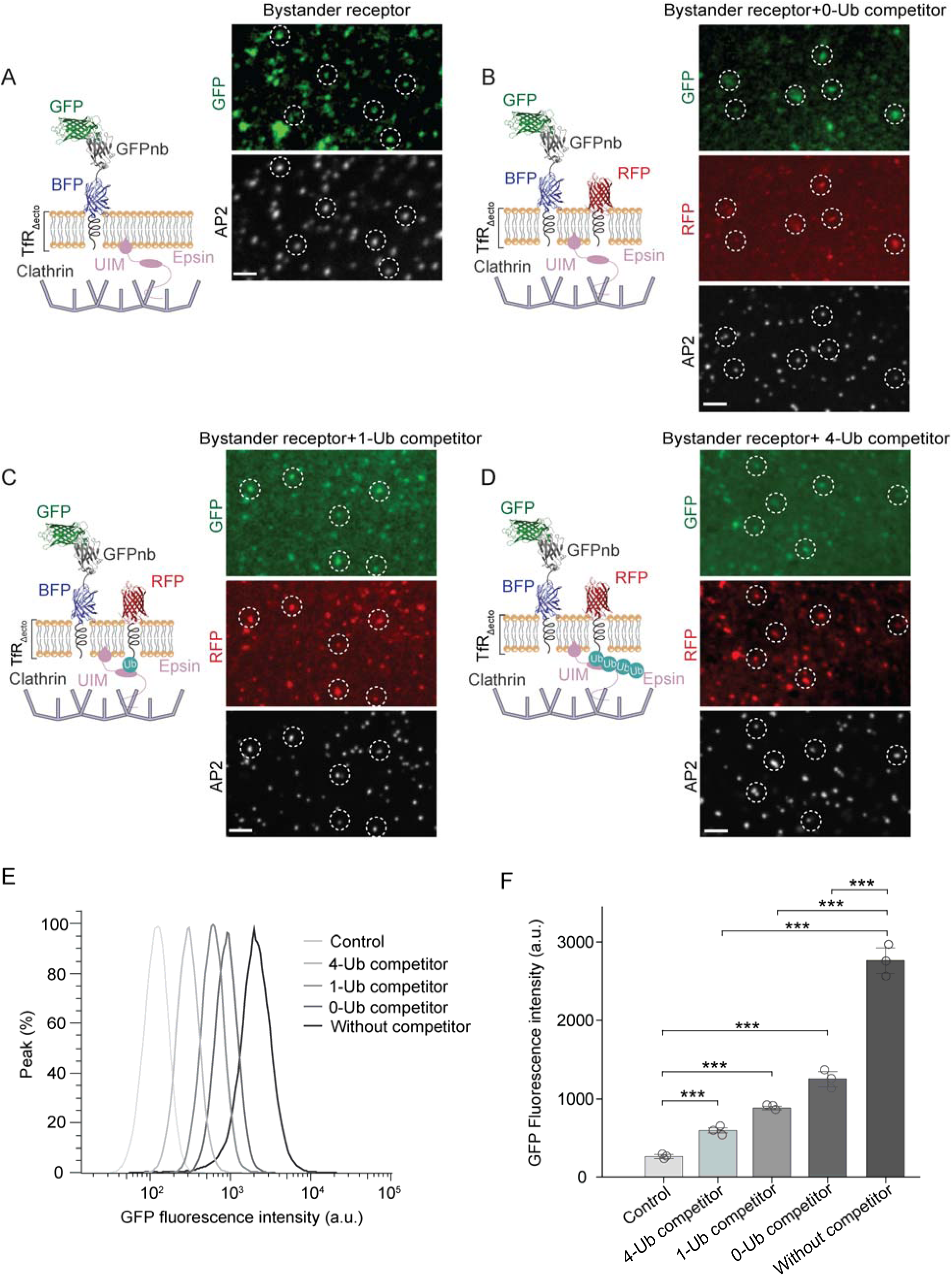
Poly-ubiquitin chain length controls ligand uptake by model receptors. (A-D) Schematic illustrations and fluorescence images showing the competition between non-ubiquitylated bystander receptors and ubiquitylated competitors for incorporation into endocytic sites. Bystander receptors (BFP-labeled) and competitors (RFP-labeled) with defined ubiquitin modifications were co-expressed in the same cells. Bystander receptor with GFPnb was used to recruit GFP for uptake. The conditions include bystander receptors expressed alone (A) or in competition with competitors modified with 0-Ub (B), 1-Ub (C), or 4-Ub(D). All lysine residues in the cytoplasmic domain of TfR were mutated to arginines to prevent ubiquitylation. To stabilize the attached ubiquitin moieties, lysines within each ubiquitin were mutated to arginines, and the C-terminal GG residues were mutated to AA to prevent intracellular deubiquitylation. Epsin binds to ubiquitylated cargo via its ubiquitin-interacting motif (UIM) and promotes clathrin-mediated endocytosis by interacting with clathrin. Fluorescence images show green fluorescence (GFP) marking non-ubiquitylated bystander receptors, red fluorescence (RFP) marking ubiquitylated competitors, and white fluorescence (AP2-JF646) indicating endocytic sites. Dashed circles highlight areas of signal colocalization between GFP, RFP, and AP2, indicating the recruitment of bystander receptors and competitors to the same endocytic sites. Scale bar: 2 µm. (E) Flow cytometry histograms showing GFP uptake across different conditions. The intensity of GFP fluorescence is higher when bystander receptors are expressed alone, while competitive incorporation into endocytic sites is reduced in the presence of competitors. Competitors with increasing levels of ubiquitylation (0-Ub, 1-Ub, and 4-Ub) progressively inhibit bystander receptors uptake, as evidenced by the leftward shift of the fluorescence peaks. (F) Quantification of GFP fluorescence intensity in model receptors under the same conditions. The GFP intensity decreases with increasing ubiquitylation levels of the competitors, demonstrating that higher ubiquitylated receptors enhances their competitive advantage for endocytic site occupancy. Data are mean ± s.e., calculated from n=3 independent biological replicates. A two-sample *t* test was conducted on the median value of the peak. P-values were < 0.05 suggesting a statistically significant difference. ***P < 0.001.

After incubation with GFP, as shown in Fig. 5A-D, we observed clear colocalization of RFP and GFP signals under confocal microscopy at the endocytic structures (AP2, JF_646_). In the RFP channel, the contrast of the endocytic sites increased with the number of ubiquitins in the chain, consistent with the results from Fig. 1. However, the contrast of GFP puncta appeared to decrease, indicating inhibition of GFP uptake in the presence of ubiquitylated competitors. Subsequently, we quantified GFP uptake using flow cytometry (*SI Appendix*, Fig. S5 and S6). Cells were incubated with GFP at 37 °C for one hour, then washed with trypsin and PBS to remove unbound GFP. As shown by the flow cytometry analysis results in Fig. 5E and F, cells expressing the bystander receptors with GFPnb exhibited a significant rightward shift in GFP fluorescence intensity distribution compared to the untransfected control group, indicating specific uptake of GFP. Notably, as the number of ubiquitin moieties on the competing receptor increased, the extent of this rightward shift in GFP fluorescence decreased significantly, Fig. 5E, F. Cells expressing the competitor with 1-Ub showed a 1.4-fold reduction in GFP uptake compared to cells expressing the 0-Ub competitor, while cells expressing the 4-Ub competitor exhibited a 2.1-fold reduction. These results collectively demonstrate the critical role of poly-ubiquitylation in mediating competition among receptors for the limited capacity of endocytic sites. In particular, the progressive reductions in uptake of non-ubiquitylated receptors in the presence of the 1-Ub and 4-Ub competitors demonstrate that cells prioritize ubiquitylated proteins for uptake at the direct expense of non-ubiquitylated proteins.

## Discussion

While ubiquitylation is thought to play a critical role in regulating the recycling of transmembrane proteins, important questions remain regarding the specific types of ubiquitylation required to promote endocytosis (11, 15, 57). In particular, it has remained unclear whether mono-ubiquitylation versus poly-ubiquitylation is sufficient to prioritize transmembrane proteins for internalization (9, 11, 26, 40). Here we demonstrate that both mono-ubiquitylation and poly-ubiquitylation can significantly enhance endocytic uptake of transmembrane proteins. However, poly-ubiquitylation had a substantially greater impact on the stability of endocytic structures in comparison to mono-ubiquitylation (Fig. 3D, E), and also conferred a larger advantage in assays of competitive localization to endocytic sites (Fig. 4) and ligand uptake (Fig. 5). Collectively, these results suggest that conjugation of poly-ubiquitin chains to transmembrane proteins plays an important role in clathrin-mediated endocytosis, both in terms of stabilizing nascent endocytic sites and prioritizing specific cargo and their ligands for removal from the plasma membrane surface.

More broadly, our findings have implications for the productive assembly of endocytic sites. It has recently been proposed that multi-valent, liquid-like assembly of endocytic proteins promotes the assembly of productive endocytic vesicles (60, 61). In particular, binding of ubiquitin to the early endocytic protein, Eps15, has been shown to shift the balance of endocytic events from abortive to productive (16). These findings are in line with other recent studies demonstrating that ubiquitin can stabilize liquid-liquid phase separation (LLPS) in diverse contexts (62–66). Our current work suggests that ubiquitylated transmembrane proteins could play an important role in stabilizing nascent endocytic sites, as suggested by data in Figure 3. In this context, poly-ubiquitylation appears to exert a more potent stabilizing effect compared to mono-ubiquitylation, suggesting that the presence of poly-ubiquitylated transmembrane cargo could promote more efficient recycling at the plasma membrane. Further, other studies suggest that ubiquitin E3 ligases of the Nedd4 family could localize within endocytic sites, where they could facilitate localized ubiquitylation of transmembrane proteins (67, 68). This effect would be expected to further stabilize endocytic sites and prioritize uptake of locally ubiquitylated cargo.

Importantly, our studies highlight the ability of poly-ubiquitylated transmembrane proteins to out-compete their non-ubiquitylated and mono-ubiquitylated counterparts during endocytic internalization. While the relationship between ubiquitylation and protein state remains poorly understood, recent work suggests that ubiquitin ligases play an important role in plasma membrane quality control, selectively targeting misfolded or damaged membrane proteins for ubiquitin-mediated internalization and degradation (69). This mechanism could help to efficiently remove defective proteins from the plasma membrane, preventing their accumulation under proteotoxic stress or after exceeding their functional lifespan. From this perspective, the ability of endocytic sites to selectively internalize poly-ubiquitylated transmembrane proteins, while leaving proteins with little or no ubiquitin behind, could provide cells with a mechanism for removing damaged proteins from the cell surface while preserving functional proteins.

## Materials and Methods

### Cell culture and transfection

Human-derived SUM159 cells gene-edited to add a HaloTag to both alleles of AP2-_σ_2 were a gift from T. Kirchhausen. Cells were grown in 1:1 DMEM high glucose: Ham’s F-12 (Hyclone, GE Healthcare) supplemented with 5% fetal bovine serum (Hyclone), Penicillin/Streptomycin/l-glutamine (Hyclone), 1 μg ml−1 hydrocortisone (H4001; Sigma-Aldrich), 5 μg ml^−1^ insulin (I6634; Sigma-Aldrich) and 10 mM HEPES, pH 7.4 and incubated at 37 °C with 5% CO_2_. Cells were seeded onto acid-washed coverslips to ensure surface cleanliness and cell adherence at a density of 3 × 10^4^ cells per coverslip for 24 h before transfection with 1 μg of plasmid DNA using 3 μL Fugene HD transfection reagent (Promega). HaloTagged AP2-_σ_2 was visualized by adding the JF_646_-HaloTag ligand (Promega). Ligand (100 nM) was added to cells and incubated at 37 °C for 15 min. Cells were washed with fresh medium and imaged immediately.

### Fluorescence microscopy

Live cell plasma membrane imaging was conducted using an Olympus spinning disk confocal microscope, which featured a Yokogawa CSU-W1 SoRa confocal scanner unit and an Olympus IX83 microscope equipped with a 100×Plan-Apochromat 1.5 NA oil-immersion objective. Fluorescence emission was captured using a Hamamatsu ORCA C13440-20CU CMOS camera. Excitation was achieved with lasers at 561 nm for RFP, and 640 nm for JF_646_.

Cell movies were collected on a TIRF microscope consisting of an Olympus IX73 microscope body, a Photometrics Evolve Delta EMCCD camera, and an Olympus 1.4 NA ×100 Plan-Apo oil objective, using Micro Manager version 1.4.23. The coverslip was heated to produce a sample temperature of 37 °C using an aluminum plate fixed to the back of the sample. All live-cell imaging was conducted in TIRF mode at the plasma membrane 24 h after transfection. Transfection media used for imaging lacked pH indicator (phenol red) and was supplemented with 1 μL OxyFluor (Oxyrase, Mansfield, OH) per 33 μL media to decrease photobleaching during live-cell fluorescence imaging. 532 nm and 640 nm lasers were used for excitation of RFP and JF_646_-HaloTag ligands of AP2, respectively. Cell movies were collected over 250 frames at 2 s intervals.

### Flow cytometry

Flow cytometry assays were conducted using a Guava easyCyte Flow Cytometer (Millipore Sigma-Aldrich), equipped with 405 nm, 488 nm, and 532 nm lasers for fluorescence excitation and detection. Samples were prepared by resuspending cells in phosphate-buffered saline (PBS) to minimize cell clumping and non-specific binding. Before acquisition, the instrument was calibrated according to the manufacturer’s recommendations to ensure consistent performance and accuracy. Data acquisition was performed at a flow rate of 35 μL/min, with event thresholds set to minimize background noise. Forward scatter (FSC) and side scatter (SSC) parameters were used to assess cell size and granularity, respectively, and gates were drawn on FSC versus SSC plots to exclude debris and identify the population of interest. Appropriate fluorescence compensation was applied to correct for spectral overlap between channels when necessary. Flow cytometry data were analyzed using FlowJo software (Treestar). Statistical analysis of flow cytometry results was performed using at least n=3 biological replicates.

### Image analysis

Clathrin-coated structures, observed as fluorescent puncta in confocal and TIRF images, were identified using CMEAnalysis (Danuser lab) (53). AP2-_σ_2 signal was used as the master channel to track clathrin-coated structures. Local intensity maxima in the RFP channel were fitted with 2D Gaussian functions. The Gaussian standard deviation was derived from the microscope’s physical parameters to estimate the point spread function. To ensure the accuracy of the fit, the Anderson-Darling test was applied to the residuals. The Gaussian amplitude, corresponding to the fluorescence intensity of each detected punctum, along with the punctum’s location, were recorded. For a punctum to qualify as a valid clathrin-coated structure, it had to be diffraction-limited and significantly brighter than the surrounding membrane, as described in previous work (36). For validated puncta in the master channel, a corresponding 2D Gaussian fit was applied to the puncta in transmembrane fusion protein, receptor, or ligand channels, within a 3_σ_ pixel radius of the master channel location (70).

Fluorescence lifetime measurements were performed by detecting and tracking clathrin-coated structures using CMEAnalysis in MATLAB. The point spread function (PSF) of the data was utilized to calculate the standard deviation of the Gaussian fit. The AP2-_σ_2 signal was designated as the master channel for tracking clathrin-coated structures. Only structures that remained visible for at least three consecutive frames were included in the analysis.

### Statistical analysis

Micrograph acquisition experiments were independently repeated on separate days, producing consistent results across replicates. For CMEAnalysis, cell image data were collected independently under at least two different experimental conditions or on separate days. As described throughout this paper, all reported experimental results are based on a minimum of three independent replicates. For comparisons with statistically significant differences, a two-sample Student’s *t*-test assuming unequal variances was used. The resulting p-values are provided in the corresponding figure captions where such comparisons are presented.

## Data, Materials and Software Availability

All study data are included in the article and/or *SI Appendix*.

## Acknowledgements

We thank T. Kirchhausen for the gift of SUM159/AP2-σ2 Halo Tag cells. This research was supported by the National Institutes of Health through grants R35GM139531 to J.C.S., the Welch Foundation through grant F-2047 to J.C.S..

## Author contributions

H.L. and J.C.S. designed the experiments. F.Y., S.S., and H.L. designed the plasmids. H.Y. performed live-cell imaging experiments. H.Y., G.A., and J.C.S. contributed to data analysis. G.A., F.Y., and C.C.H. assisted with the calibration and setup of the imaging microscope. H.L. and J.C.S. wrote the manuscript, with J.M.H. and S.S. providing consultation on manuscript preparation and editing.

## References

1. Goh, L.K. & Sorkin, A. Endocytosis of receptor tyrosine kinases. Cold Spring Harb. Perspect. Biol. 5, a017459 (2013).

2. Sorkin, A. & Von Zastrow, M. Endocytosis and signalling: intertwining molecular networks. Nat. Rev. Mol. Cell Biol. 10, 609–622 (2009).

3. Vieira, A.V., Lamaze, C. & Schmid, S.L. Control of EGF receptor signaling by clathrin-mediated endocytosis. Science 274, 2086–2089 (1996).

4. McMahon, H.T. & Boucrot, E. Molecular mechanism and physiological functions of clathrin-mediated endocytosis. Nat. Rev. Mol. Cell Biol. 12, 517–533 (2011).

5. Cocucci, E., Aguet, F., Boulant, S. & Kirchhausen, T. The first five seconds in the life of a clathrin-coated pit. Cell 150, 495–507 (2012).

6. Kaksonen, M. & Roux, A. Mechanisms of clathrin-mediated endocytosis. Nat. Rev. Mol. Cell Biol. 19, 313–326 (2018).

7. Schmid, E.M. & McMahon, H.T. Integrating molecular and network biology to decode endocytosis. Nature 448, 883–888 (2007).

8. Antonny, B. et al. Membrane fission by dynamin: what we know and what we need to know. EMBO J. 35, 2270–2284 (2016).

9. Sigismund, S. et al. Clathrin-independent endocytosis of ubiquitinated cargos. Proc. Natl. Acad. Sci. U.S.A. 102, 2760–2765 (2005).

10. Haglund, K. & Dikic, I. The role of ubiquitylation in receptor endocytosis and endosomal sorting. J. Cell Sci. 125, 265–275 (2012).

11. Marmor, M.D. & Yarden, Y. Role of protein ubiquitylation in regulating endocytosis of receptor tyrosine kinases. Oncogene 23, 2057–2070 (2004).

12. Kozak, M. & Kaksonen, M. Condensation of Ede1 promotes the initiation of endocytosis. Elife 11, e72865 (2022).

13. Hicke, L. & Dunn, R. Regulation of membrane protein transport by ubiquitin and ubiquitin-binding proteins. Annu. Rev. Cell Dev. Biol. 19, 141–172 (2003).

14. Polo, S. et al. A single motif responsible for ubiquitin recognition and monoubiquitination in endocytic proteins. Nature 416, 451–455 (2002).

15. Riezman, H. The ubiquitin connection. Nature 416, 381–383 (2002).

16. Yuan, F. et al. Ubiquitin-driven protein condensation stabilizes clathrin-mediated endocytosis. PNAS Nexus 3, 342 (2024).

17. Shih, S.C. et al. Epsins and Vps27p/Hrs contain ubiquitin-binding domains that function in receptor endocytosis. Nat. Cell Biol. 4, 389–393 (2002).

18. Chen, H. & De Camilli, P. The association of epsin with ubiquitinated cargo along the endocytic pathway is negatively regulated by its interaction with clathrin. Proc. Natl. Acad. Sci. U.S.A. 102, 2766–2771 (2005).

19. Chen, H. et al. Epsin is an EH-domain-binding protein implicated in clathrin-mediated endocytosis. Nature 394, 793–797 (1998).

20. Rosenthal, J.A. et al. The epsins define a family of proteins that interact with components of the clathrin coat and contain a new protein module. J. Biol. Chem. 274, 33959–33965 (1999).

21. Wendland, B. Epsins: adaptors in endocytosis? Nat. Rev. Mol. Cell. Biol. 3, 971–977 (2002).

22. Hawryluk, M.J. et al. Epsin 1 is a polyubiquitin_-_selective clathrin_-_associated sorting protein. Traffic 7, 262–281 (2006).

23. Barriere, H. et al. Molecular basis of oligoubiquitin_-_dependent internalization of membrane proteins in Mammalian cells. Traffic 7, 282–297 (2006).

24. Traub, L.M. Tickets to ride: selecting cargo for clathrin-regulated internalization. Nat. Rev. Mol. Cell. Biol. 10, 583–596 (2009).

25. Hochstrasser, M. Ubiquitin-dependent protein degradation. Annu. Rev. Genet. 30, 405–439 (1996).

26. Hicke, L. Protein regulation by monoubiquitin. Nat. Rev. Mol. Cell. Biol. 2, 195–201 (2001).

27. Galan J M, Haguenauer_-_Tsapis R. Ubiquitin lys63 is involved in ubiquitination of a yeast plasma membrane protein. EMBO J. 16, 5847–5854 (1997).

28. Galan J M, Moreau V, Andre B, et al. Ubiquitination Mediated by the Npi1p/Rsp5p Ubiquitin-protein Ligase Is Required for Endocytosis of the Yeast Uracil Permease. J. Biol. Chem. 271, 10946–10952 (1996).

29. Huang, F., Kirkpatrick, D., Jiang, X., Gygi, S. & Sorkin, A. Differential regulation of EGF receptor internalization and degradation by multiubiquitination within the kinase domain. Mol. Cell 21, 737–748 (2006).

30. Mosesson, Y. et al. Endocytosis of receptor tyrosine kinases is driven by monoubiquitylation, not polyubiquitylation. J. Biol. Chem. 278, 21323–21326 (2003).

31. DeGroot, A.C.M., Gollapudi, S., Zhao, C., LaMonica, M.F. & Stachowiak, J.C. Weakly Internalized Receptors Use Coated Vesicle Heterogeneity to Evade Competition during Endocytosis. Biochemistry 60, 2195–2205 (2021).

32. Zhao, C. et al. Receptor Heterodimerization Modulates Endocytosis through Collaborative and Competitive Mechanisms. Biophys. J. 117, 646–658 (2019).

33. DeGroot, A.C.M., Zhao, C., LaMonica, M.F., Hayden, C.C. & Stachowiak, J.C. Molecular thermodynamics of receptor competition for endocytic uptake. Soft Matter 15, 7448–7461 (2019).

34. Marks, M.S., Woodruff, L., Ohno, H. & Bonifacino, J.S. Protein targeting by tyrosine-and di-leucine-based signals: evidence for distinct saturable components. J. Cell Biol. 135, 341–354 (1996).

35. Warren, R.A., Green, F.A., Stenberg, P.E. & Enns, C.A. Distinct saturable pathways for the endocytosis of different tyrosine motifs. J. Biol. Chem. 273, 17056–17063 (1998).

36. Gollapudi, S. et al. Steric pressure between glycosylated transmembrane proteins inhibits internalization by endocytosis. Proc. Natl. Acad. Sci. U.S.A. 120, e2215815120 (2023).

37. Lauffenburger, D.A. & Linderman, J.J. Receptors: models for binding, trafficking, and signaling. (Oxford University Press, USA, 1996).

38. Voss, P. & Grune, T. The nuclear proteasome and the degradation of oxidatively damaged proteins. Amino acids 32, 527–534 (2007).

39. Pajares, M. et al. Redox control of protein degradation. Redox Biol. 6, 409–420 (2015).

40. Piper, R.C., Dikic, I. & Lukacs, G.L. Ubiquitin-dependent sorting in endocytosis. Cold Spring Harb. Perspect. Biol. 6, a016808 (2014).

41. Harding, C., Heuser, J. & Stahl, P. Receptor-mediated endocytosis of transferrin and recycling of the transferrin receptor in rat reticulocytes. J. Cell Biol. 97, 329–339 (1983).

42. Collawn, J.F. et al. Transferrin receptor internalization sequence YXRF implicates a tight turn as the structural recognition motif for endocytosis. Cell 63, 1061–1072 (1990).

43. Collawn, J. et al. YTRF is the conserved internalization signal of the transferrin receptor, and a second YTRF signal at position 31-34 enhances endocytosis. J. Biol. Chem. 268, 21686–21692 (1993).

44. Ohno, H. et al. Interaction of tyrosine-based sorting signals with clathrin-associated proteins. Science 269, 1872–1875 (1995).

45. Kelly, B.T. et al. A structural explanation for the binding of endocytic dileucine motifs by the AP2 complex. Nature 456, 976–979 (2008).

46. Sandoval, I.V. & Bakke, O. Targeting of membrane proteins to endosomes and lysosomes. Trends Cell Biol. 4, 292–297 (1994).

47. Géminard, C., De Gassart, A., Blanc, L. & Vidal, M. Degradation of AP2 during reticulocyte maturation enhances binding of hsc70 and Alix to a common site on TFR for sorting into exosomes. Traffic 5, 181–193 (2004).

48. Fujita, H., Iwabu, Y., Tokunaga, K. & Tanaka, Y. Membrane-associated RING-CH (MARCH) 8 mediates the ubiquitination and lysosomal degradation of the transferrin receptor. J. Cell Sci. 126, 2798–2809 (2013).

49. Collins, B.M., McCoy, A.J., Kent, H.M., Evans, P.R. & Owen, D.J. Molecular architecture and functional model of the endocytic AP2 complex. Cell 109, 523–535 (2002).

50. Owen, D.J., Collins, B.M. & Evans, P.R. Adaptors for clathrin coats: structure and function. Annu. Rev. Cell Dev. Biol. 20, 153–191 (2004).

51. Blondeau, F. et al. Tandem MS analysis of brain clathrin-coated vesicles reveals their critical involvement in synaptic vesicle recycling. Proc. Natl. Acad. Sci. USA 101, 3833–3838 (2004).

52. Grimm, J.B. et al. A general method to improve fluorophores for live-cell and single-molecule microscopy. Nat. Methods 12, 244–250 (2015).

53. Aguet, F., Antonescu, C.N., Mettlen, M., Schmid, S.L. & Danuser, G. Advances in analysis of low signal-to-noise images link dynamin and AP2 to the functions of an endocytic checkpoint. Dev. Cell 26, 279–291 (2013).

54. DeGroot, A.C.M. et al. Entropic Control of Receptor Recycling Using Engineered Ligands. Biophys. J. 114, 1377–1388 (2018).

55. Kubala, M.H., Kovtun, O., Alexandrov, K. & Collins, B.M. Structural and thermodynamic analysis of the GFP: GFP_-_nanobody complex. Protein Sci. 19, 2389–2401 (2010).

56. Gadok, A.K. et al. Connectosomes for Direct Molecular Delivery to the Cellular Cytoplasm. J. Am. Chem. Soc. 138, 12833–12840 (2016).

57. Weinberg, J.S. & Drubin, D.G. Regulation of clathrin-mediated endocytosis by dynamic ubiquitination and deubiquitination. Curr. Biol. 24, 951–959 (2014).

58. Loerke, D. et al. Cargo and dynamin regulate clathrin-coated pit maturation. PLoS Biol. 7, e1000057 (2009).

59. Boulant, S., Kural, C., Zeeh, J.-C., Ubelmann, F. & Kirchhausen, T. Actin dynamics counteract membrane tension during clathrin-mediated endocytosis. Nat. Cell Biol. 13, 1124–1131 (2011).

60. Day, K.J. et al. Liquid-like protein interactions catalyse assembly of endocytic vesicles. Nat. Cell Biol. 23, 366–376 (2021).

61. Huang, W.Y. et al. Phosphotyrosine-mediated LAT assembly on membranes drives kinetic bifurcation in recruitment dynamics of the Ras activator SOS. Proc. Natl. Acad. Sci. USA 113, 8218–8223 (2016).

62. Du, Z. et al. DNMT1 stability is regulated by proteins coordinating deubiquitination and acetylation-driven ubiquitination. Sci. Signal. 3, 146 (2010).

63. Stack, J.H., Whitney, M., Rodems, S.M. & Pollok, B.A. A ubiquitin-based tagging system for controlled modulation of protein stability. Nat. Biotechnol. 18, 1298–1302 (2000).

64. Lee, J. & Gu, W. The multiple levels of regulation by p53 ubiquitination. Cell Death Differ. 17, 86–92 (2010).

65. Popov, N. et al. The ubiquitin-specific protease USP28 is required for MYC stability. Nat. Cell Biol. 9, 765–774 (2007).

66. Li, M. et al. Deubiquitination of p53 by HAUSP is an important pathway for p53 stabilization. Nature 416, 648–653 (2002).

67. Wang, G. et al. Localization of the Rsp5p ubiquitin-protein ligase at multiple sites within the endocytic pathway. Mol. Cell. Biol. 21, 3564–3575 (2001).

68. Yang, X., Arines, F.M., Zhang, W. & Li, M. Sorting of a multi-subunit ubiquitin ligase complex in the endolysosome system. Elife 7, e33116 (2018).

69. Zhao, Y., MacGurn, J.A., Liu, M. & Emr, S. The ART-Rsp5 ubiquitin ligase network comprises a plasma membrane quality control system that protects yeast cells from proteotoxic stress. Elife 2, e00459 (2013).

70. Ashby, G., Keng, K.E., Hayden, C.C. & Stachowiak, J.C. A live cell imaging-based assay for tracking particle uptake by clathrin mediated endocytosis. BioRvix 2, 579544 (2024).

