## Supplementary Information for "Poly-ubiquitylated transmembrane proteins outcompete other cargo for limited space inside clathrin-coated vesicles"

Hao-Yang Liu *et al*.

**This PDF file includes:**

Supplementary Information

Figs. S1 to S6

References

**Supporting Materials and Methods**

**Plasmids**

A plasmid for expression of 0-Ub-TfR-Δecto-RFP was generated by replacing the GFP domain with mRFP of the TfR-Δecto-GFP construct, which we previously described (1). The TfR-Δecto-GFP plasmid encoded the first 88 amino acids of the human transferrin receptor, corresponding to the intracellular and transmembrane domains of TfR (identical to amino acids 1-88 of GenBank accession number AAA61153) fused to GFP. A 9-amino acid linking sequence (GKGDPPVAT) connected TfR and GFP in this construct. Here mRFP was PCR amplified from the pcDNA3-mRFP plasmid acquired through Addgene as a gift from Dr. Douglas Golenbock (Addgene #13032). To prevent ubiquitylation, all four lysine residues in the cytoplasmic domain of TfR were mutated to arginines (K to R) to prevent ubiquitylation.

The constructs 1-Ub-TfR-Δecto-mRFP and 4-Ub-TfR-Δecto-mRFP were generated by adding one or four tandem ubiquitin (Ub) coding sequences, respectively, to the N-terminus of the 0-Ub-TfR-Δecto-RFP plasmid. To stabilize the attached ubiquitin moieties and prevent unintended poly-ubiquitylation, all seven lysine residues within the ubiquitin sequence were mutated to arginines (K to R). Additionally, to prevent intracellular deubiquitylation, the C-terminal glycine residues (GG) of ubiquitin were mutated to alanine residues (GG to AA). This design ensured that the ubiquitin chains remained covalently attached and stable during cellular expression. For the 1-Ub-TfR-Δecto-RFP construct, a single mutated ubiquitin sequence was added to the N-terminus of TfR-Δecto-RFP. For the 4-Ub-TfR-Δecto-RFP construct, four tandemly fused ubiquitin sequences with the same mutations were inserted at the same position.

To generate 0-Ub-TfR-Δecto-mRFP-GFPnb, 1-Ub-TfR-Δecto-mRFP-GFPnb, and 4-Ub-TfR-Δecto-mRFP-GFPnb, the GFP nanobody (GFPnb) coding sequence was PCR amplified from the pOPINE GFP nanobody plasmid (Addgene #49172), a gift from Dr. Brett Collins. The amplified GFPnb sequence was inserted into the C-terminus of each plasmid, downstream of the mRFP coding region. To ensure flexibility and proper folding of the fused proteins, a flexible linker sequence (ESRPQYNQRRYT) was introduced between the mRFP and GFPnb coding regions. The resulting constructs were designed to retain the functional properties of both mRFP and GFPnb while allowing proper spatial arrangement for nanobody binding to GFP-tagged proteins.

To generate the constructs 0-Ub-TfR-Δecto-GFP, 1-Ub-TfR-Δecto-GFP, and 4-Ub-TfR-Δecto-GFP, the mRFP coding region in the respective 0-Ub-TfR-Δecto-RFP, 1-Ub-TfR-Δecto-RFP, and 4-Ub-TfR-Δecto-RFP plasmids was replaced with the coding sequence for GFP. To improve the monomeric properties of GFP and prevent dimerization, the A206K mutation was introduced into the GFP sequence using site-directed mutagenesis. This mutation converts alanine at position 206 to lysine, ensuring that the GFP remains monomeric under physiological conditions.

To generate the 0-Ub-TfR-Δecto-BFP-GFPnb construct, all four lysine residues in the cytoplasmic domain of TfR were mutated to arginines (K to R) to prevent ubiquitylation. The TfR-Δecto-BFP-GFPnb plasmid, which we had previously used (2), served as the template for this modification.

All plasmid constructs above were custom-designed and constructed by GenScript. The designs included specific modifications such as the addition of ubiquitin sequences with lysine-to-arginine (K to R) mutations and glycine-to-alanine (GG to AA) modifications, replacement of mRFP with GFP (A206K mutant) or BFP, and the inclusion of GFP nanobody (GFPnb) with a flexible linker. After receiving the plasmids, all constructs were verified by full sequencing (Plasmidsaurus) to confirm the correct insertion, reading frame, and presence of the intended mutations.

**Supplementary Figures**

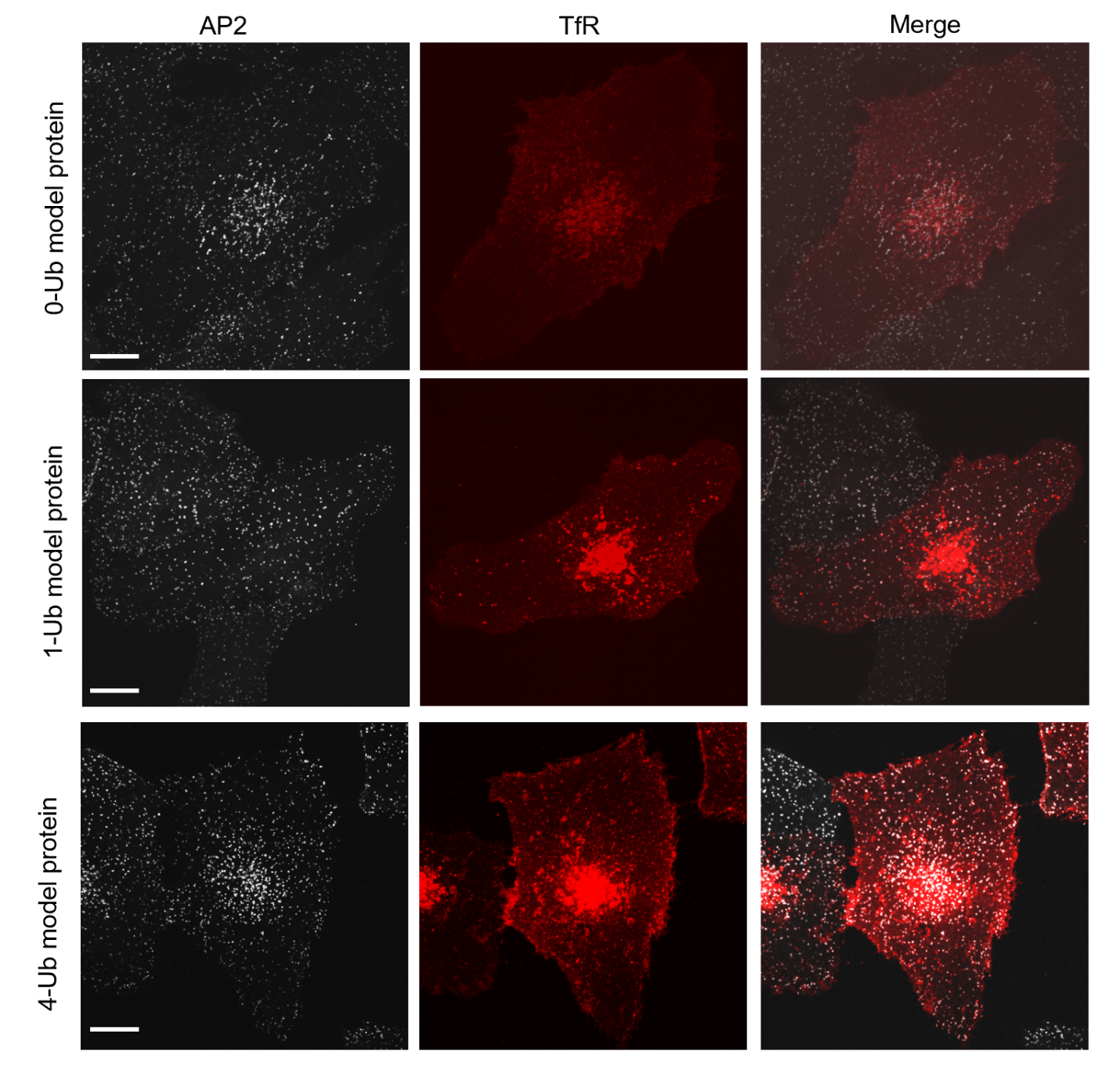

**Fig. S1**. Whole cell fluorescence images of the plasma membrane of SUM 159 cells expressing the model proteins. (A-C) Spinning disk confocal images of whole cells expressing the 0-Ub (A), 1-Ub (B) and 4-Ub model proteins. Crops of these cells are shown in Figure 1 D-F, respectively. Red fluorescence (RFP) highlights the model proteins, while white fluorescence (AP2-JF_646_) marks endocytic sites. The scale bars are 10 µm.

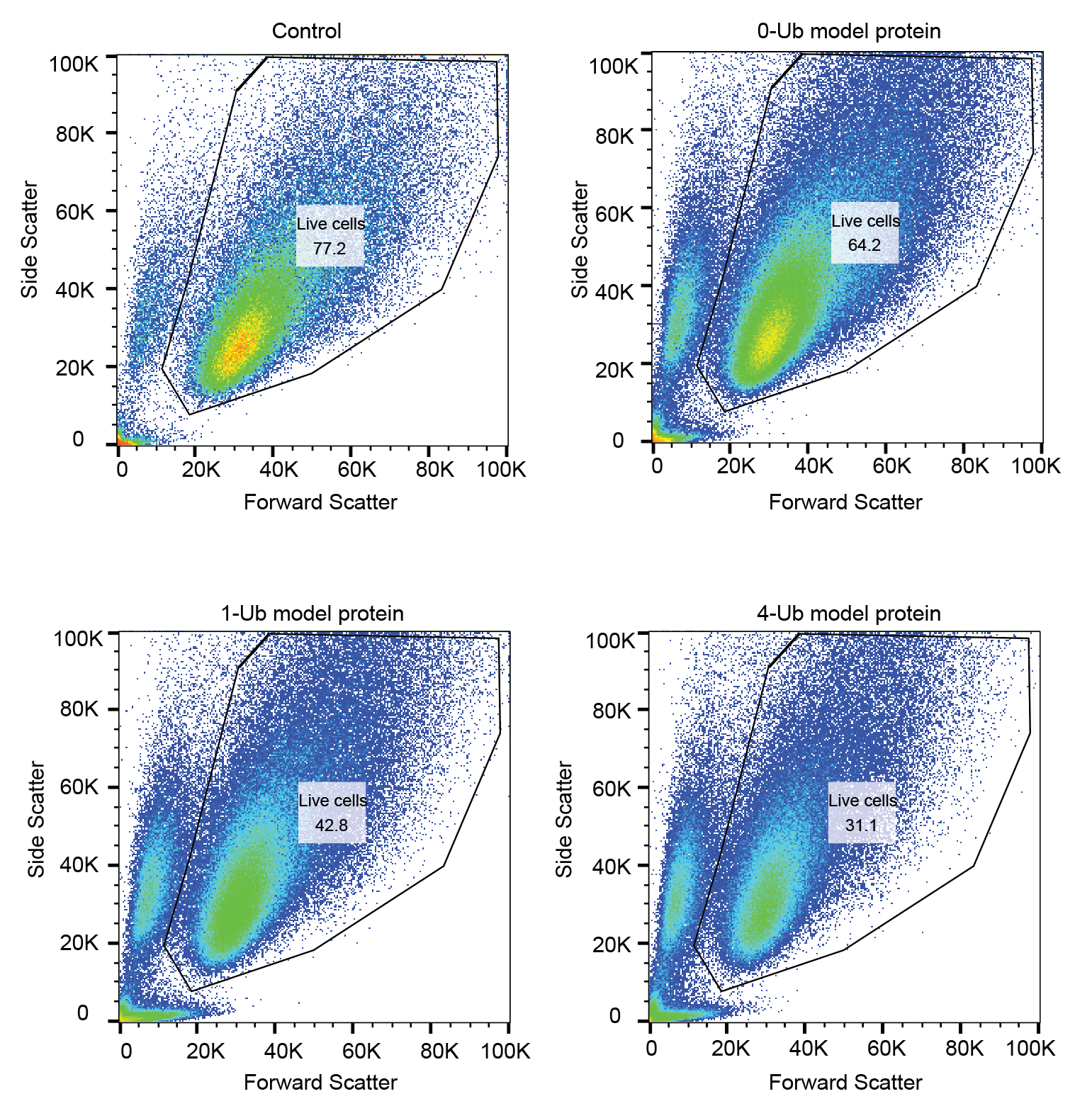

**Fig. S2**. Flow cytometry scatter plots from populations of cells of each condition in Figure 2D. The black irregular circle represents the gate. n=3 trials were done for each condition.

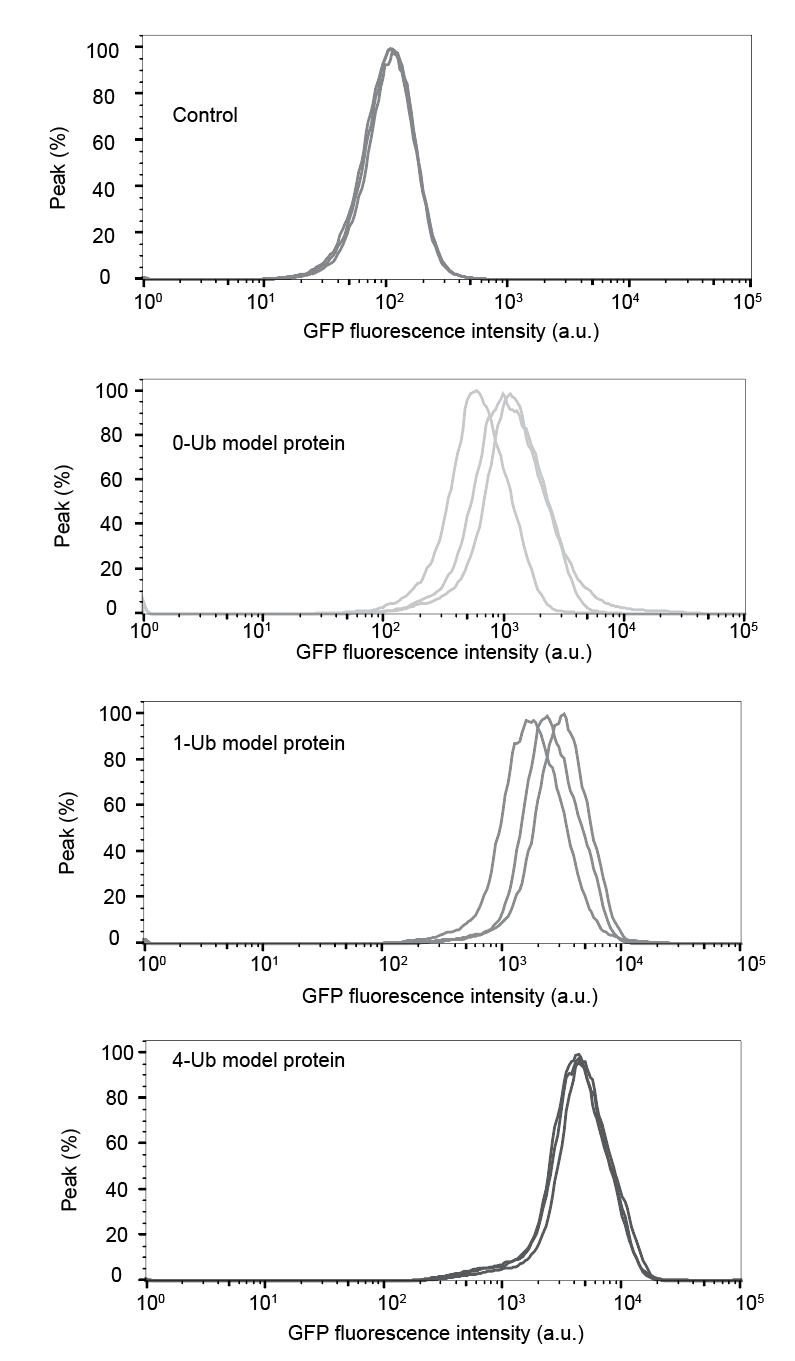

**Fig. S3**. Flow cytometry histograms of the GFP fluorescence intensity of cells under each condition presented in Figure 2D. Each condition was assessed in three independent trials (n=3).

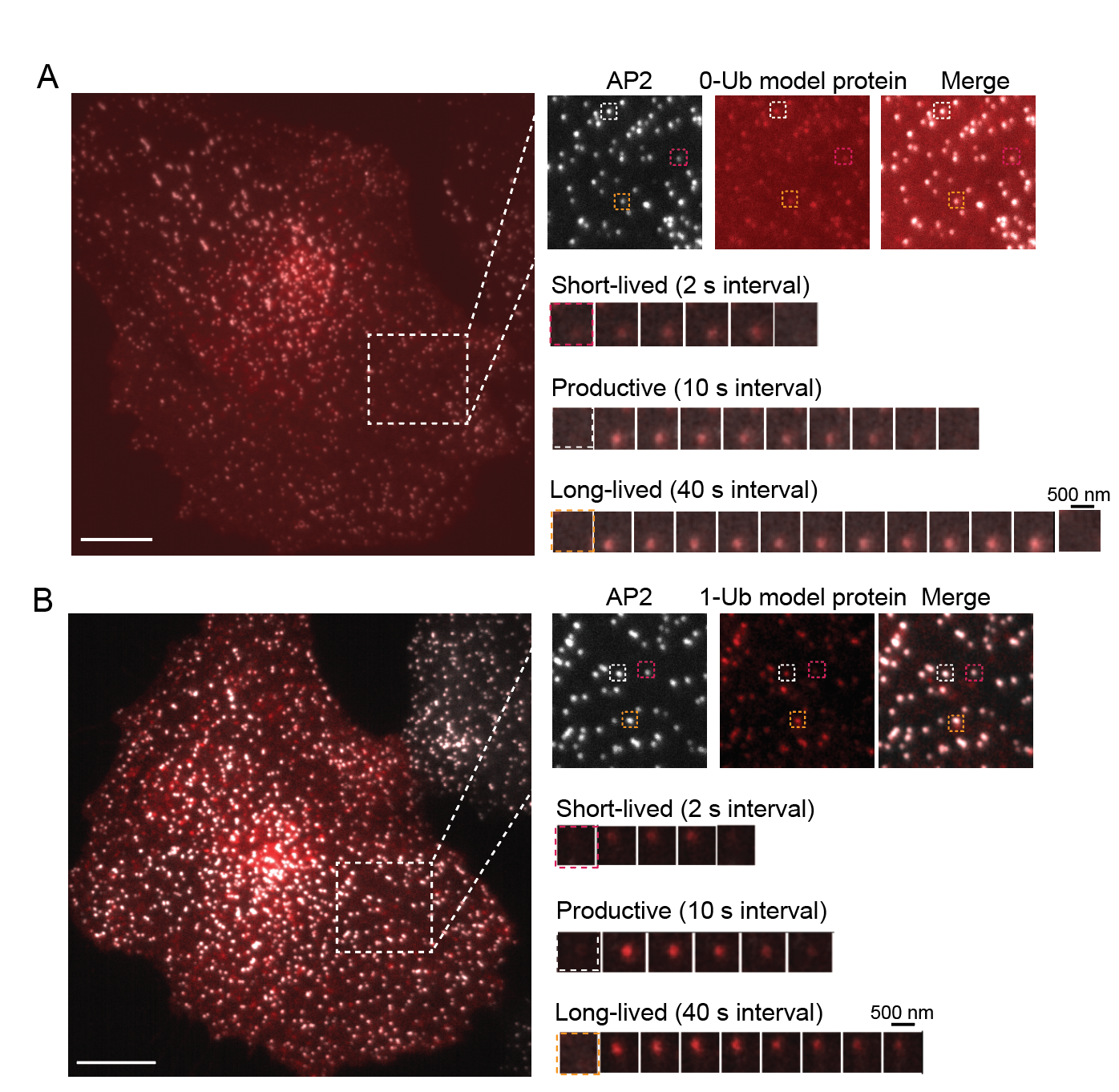

**Fig. S4**. Representative TIRF images of SUM159 cells expressing 0-Ub (A) and 1-Ub (B) model protein: AP2 (white) and 4-Ub model protein (red). Large inset highlights three representative clathrin-coated structures shown in smaller insets: short-lived (red), productive (white), and long-lived (orange) structures, respectively. Scale bars: 10 µm (main image), 500 nm (insets).

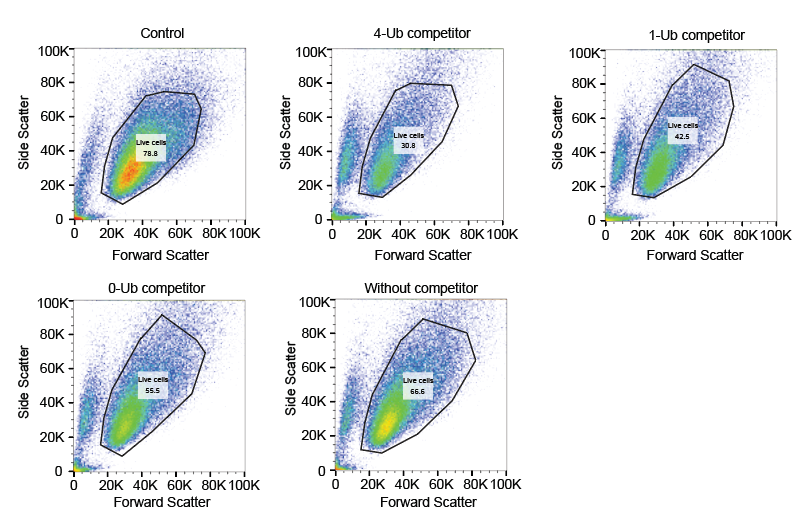

**Fig. S5**. Flow cytometry scatter plots from populations of cells of each condition in Figure 5E. The black irregular circle represents the gate. n=3 trials were done for each condition.

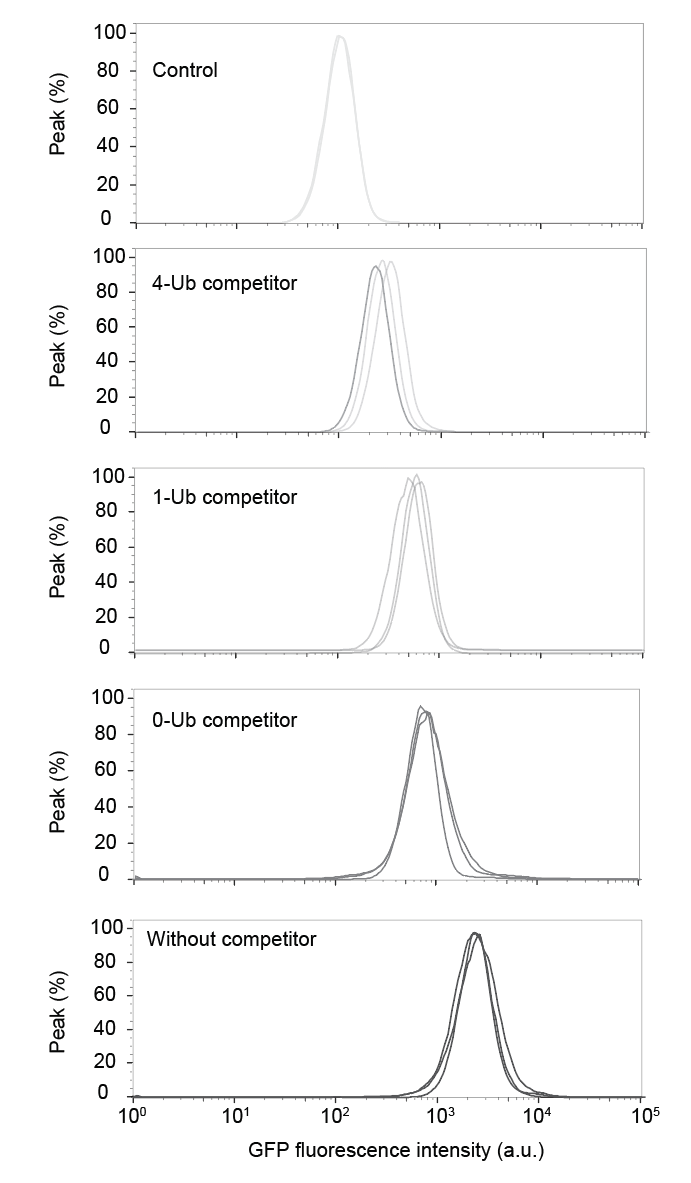

**Fig. S6**. Flow cytometry histograms of the GFP fluorescence intensity of cells under each condition presented in Figure 5E. Each condition was assessed in three independent trials (n=3).

**Reference:**

1. Busch, D.J., J.R. Houser, C.C. Hayden, M.B. Sherman, E.M. Lafer, and J.C. Stachowiak. Intrinsically disordered proteins drive membrane curvature. *Nat. Commun*. **6**, 7875 (2015).
2. DeGroot, A.C.M. et al. Entropic Control of Receptor Recycling Using Engineered Ligands. *Biophys. J*. **114**, 1377-1388 (2018).
